# THIK-1 channel mediates microglial glucose sensing and modulates AgRP neurons

**DOI:** 10.64898/2026.01.15.699765

**Authors:** Qingzhuo Liu, Jonathan C. Bean, Jinjing Jian, Tong Zhou, Yuxue Yang, Meixin Sun, Yongxiang Li, Kristine M. Conde, Mengjie Wang, Yue Deng, Jiamin Qiu, Fuhui Wang, Xinming Liu, Yutian Liu, Jingjing Cheng, Xi Wu, Lamei Xue, Chunling Chen, Chenling Meng, Yi Zhu, Yongjie Yang, Longlong Tu, Hailan Liu, Yong Xu

## Abstract

Microglia play essential roles in maintaining energy homeostasis, and their dysfunction contributes to metabolic disease. Although high-fat diet (HFD) exposure induces microglial activation, the underlying mechanisms remain poorly defined. Here, we identified a previously unrecognized role for THIK-1 channel in mediating glucose sensing of microglia in arcuate nucleus of the hypothalamus (ARH), during HFD-induced obesity. Pharmacological inhibition of THIK-1 channel with tetrapentylammonium (TPA) suppresses feeding and attenuates body-weight gain in diet induced obese mice. Mechanistically, inhibition of agouti-related peptide (AgRP) neurons is indispensable for TPA-induced hypophagia. Moreover, THIK-1 inhibition promotes microglial phagocytosis of perineuronal nets (PNNs), leading to reduced AgRP neuronal activity and feeding suppression. Together, these findings establish THIK-1 as a critical glucose sensor in hypothalamic microglia and uncover a microglia-dependent pathway through which overnutrition modulates AgRP neuronal activity via PNN remodeling to regulate energy balance, highlighting THIK-1 as a potential therapeutic target for treatment of diet-induced obesity.

## INTRODUCTION

The prevalence of obesity is increasing worldwide^1^. This global epidemic continues to advance unabated and currently affects more than two billion individuals, representing approximately 30% of the world’s population^2^. The rising incidence of obesity and overweight is accompanied by a substantial increase in associated health complications, including insulin resistance, type 2 diabetes, cardiovascular disease, liver disease, cancer, and neurodegenerative disorders^3–5^. Caloric intake and energy expenditure contribute to body weight regulation, accumulating evidence indicates that overeating plays a more prominent role in the development of obesity than reductions in physical activity^6^. In particular, chronic exposure to highly palatable, energy-dense foods may override homeostatic metabolic demands and cause a loss of control in the regulation of food intake^1^. While this framework provides an appealing explanation for the obesity epidemic, the neurobiological mechanisms underlying such aberrant feeding behavior remain poorly understood. Thus, it is of paramount importance to elucidate how pathological overeating emerges, both to advance mechanistic understanding and to identify novel targets for therapeutic intervention.

Glucose homeostasis requires that organism rapidly responds to fluctuations in glucose levels in the body^7^. Given that the central nervous system depends on a continuous supply of glucose, it needs to detect changes in glucose availability and coordinate with peripheral organ function to maintain its energetic demands^8^. Nearly half a century ago, specialized glucose-sensing neurons were identified as key components of this regulatory system^8^. These neurons use glucose as the metabolic signal and adjust their firing rates accordingly: glucose-excited neurons (GE neurons) increase activity as glucose levels rise, whereas glucose-inhibited neurons (GI neurons) fire more rapidly when glucose levels fall and reduce their activity as glucose levels increase^8^. Such glucose-sensing neurons are distributed across several brain regions, including the ventromedial hypothalamic nucleus (VMH), the arcuate nucleus of hypothalamus (ARH), the paraventricular nucleus of the hypothalamus (PVH), the nucleus of the solitary tract (NTS), and the medial amygdala (MeA)^9–12^. In addition to these glucose-sensing neurons, astrocytes play a pivotal role in brain energy metabolism by mediating glucose uptake, sensing metabolic fluctuations, and modulating synaptic activity^13^. Specifically, hypothalamic astrocytes adjust their lactate production in response to glucose fluctuations in an AMPK (AMP-activated protein kinase)-dependent manner, establishing themselves as glucose responsive cells^14^. Meanwhile, microglia perform essential homeostatic functions in the brain, and their pronounced plasticity and responsiveness to environmental cues allow them to rapidly transition into non-homeostatic states^15^. For example, microglia in the mediobasal hypothalamus (MBH) act as sensors of rising dietary saturated fatty acids and regulate a highly localized inflammatory response^16^. Moreover, microglial inflammatory signaling has been implicated in central regulation of glucose homeostasis; activating microglia improves glucose tolerance in high-fat diet (HFD)-fed mice in a manner linked to modulation of hypothalamic glucose-sensing neurons^15^. However, whether microglia themselves directly detect changes in glucose availability remains unknown.

Microglia are resident immune cells in the central nervous system that are intimately involved in both physiological and pathological processes in the brain^15,17,18^. Through highly motile fine processes, microglia continuously survey the tissue microenvironment and become rapidly activated in response to subtle alterations in synaptic activity as well as pathogen-associated molecular patterns^19,20^. In the hypothalamus, microglia have been proposed to function as local sensors capable of orchestrating region-specific inflammatory responses^21^. Consistent with this role, hypothalamic inflammation has been shown to disrupt energy balance regulation by altering hypothalamic circuitry^22,23^. For example, hypercaloric diets stimulate microglial Toll-like receptor 4 (TLR4) signaling and induce tumor necrosis factor-α (TNFα) secretion, leading to increased excitability of pro-opiomelanocortin (POMC) neurons in the ARH^24,25^. In parallel with their role in modulating neuronal activity in the adult brain, microglia are also essential for postnatal hypothalamic circuit maturation. Microglia-mediated synaptic pruning is essential for postnatal synaptic maturation, and microglial depletion during a critical postnatal period of ARH neuron development increases both the number of AgRP neurons and their fiber density, indicating that microglia actively participate in shaping AgRP neuronal circuits^26^. Although AgRP and POMC neurons mediate orexigenic and anorexigenic signaling within the ARH^27^, how microglia interact with these neuronal populations to regulate feeding behavior in the context of obesity remains incompletely understood.

In the current study, we identified THIK-1 channel as critical mediators of glucose-sensing in ARH microglia in the context of HFD induced obesity. Notably, inhibition of microglial THIK-1 channel suppresses feeding and attenuates body-weight gain in obese mice. Mechanistically, we found that inhibition of AgRP neurons is required for TPA-induced feeding suppression. Furthermore, THIK-1 inhibition promotes microglial phagocytosis of PNNs, and that PNNs remodeling represents a critical mechanism linking microglial THIK-1 inhibition to altered AgRP neuronal activity and feeding suppression. Collectively, these findings establish THIK-1 as a critical glucose-sensing channel in ARH microglia and uncover a novel microglia-dependent pathway through which THIK-1 channel regulate energy balance, highlighting it as potential therapeutic target for obesity treatment.

## RESULTS

### THIK-1 mediates glucose sensing of microglia in the ARH

Previous studies have shown that HFD feeding promotes microglial accumulation and increases microglial cell size within the ARH in both rats and mice^28,29^. To dissect the mechanism underlying this microglial response, we fed Tmem119-CreER/Rosa26-LSL-tdTomato male mice (microglia are labeled with tdTomato in this mouse line) with chow diet or HFD for 1 week and then prepared ARH-containing brain slices for microglial electrophysiological recordings (Figure 1A and 1B). We found that the one-week HFD feeding significantly hyperpolarized ARH microglia (Figure 1C and 1D). Microglial membrane potential, which is highly related to chemotactic response, has been implicated in controlling microglial ramification and surveillance^30–32^. To further investigate how HFD exposure induces microglial hyperpolarization, we next examined the resting membrane potential of ARH microglia under current-clamp mode in response to a 1 → 5 mM or 5 →1 mM extracellular glucose fluctuation protocol (see Methods), as previously described^33,34^. During euglycemia, brain glucose concentrations are maintained within a narrow range of 1–2.5 mM^35^, whereas under hyperglycemic conditions they rise to ∼5 mM^36^. Strikingly, increasing extracellular glucose from 1 mM to 5 mM significantly decreased the resting membrane potential (Figure 1E and 1F), whereas reducing glucose from 5 mM to 1 mM produced the opposite effect and depolarized microglia (Figure 1E and 1F). Together, these results demonstrated that ARH microglia are glucose-sensing.

**Figure 1.**
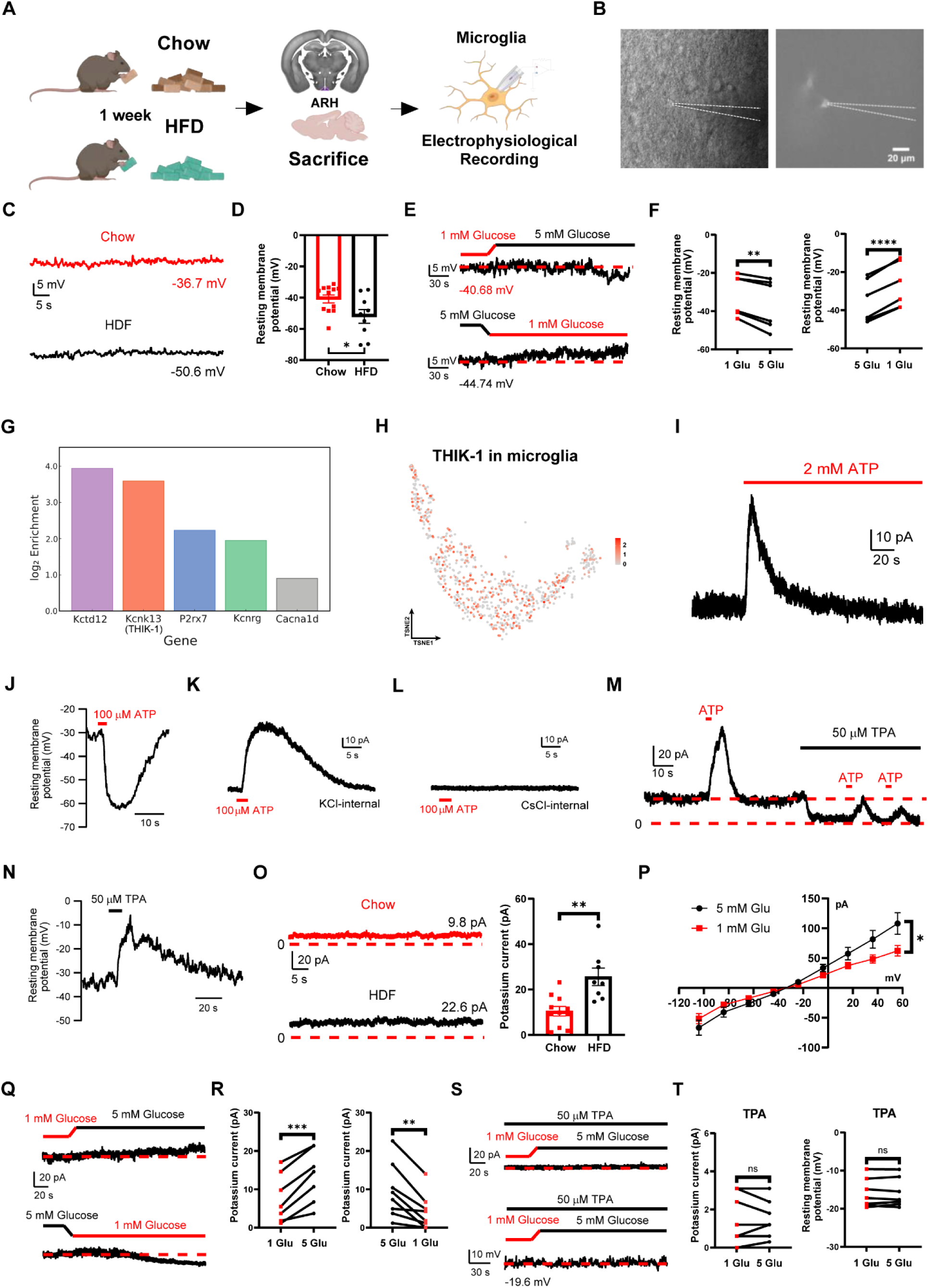
THIK-1 mediates glucose sensing of microglia in the ARH. (A) The experimental design for recording microglial activity in the ARH after chow or HFD exposure for 1 week. (B) Experimental illustration of recorded microglia. (C) Representative resting membrane potential of ARH microglia recorded after chow or HFD exposure for 1 week in male Tmem119-CreER/Rosa26-LSL-tdTomato mice. (D) Statistics for resting membrane potential of ARH microglia following chow or HFD exposure. N = 12 (chow) or 9 (HFD) microglia from 3 different mice in each group. \**p* < 0.05. (E) Representative resting membrane potential responses to glucose fluctuation (1 → 5 mM or 5 →1 mM) in ARH microglia. (F) Statistics for resting membrane potential of microglia in the ARH, under glucose exposure from 1 → 5 mM (n = 6) or 5 →1 mM (n = 6). Significant differences between groups are shown as \*\**p* < 0.01 and \*\*\*\**p* < 0.0001. (G) Bar plot showing the top five enriched ion channels in microglia of the ARH. (H) Feature plot showing THIK-1 expression in microglia of the ARH. (I) Representative voltage-clamp trace of microglial current recorded at 0 mV evoked by 2 mM ATP. (J) Representative resting membrane potential of ARH microglia response to ATP puff (100 µM). (K-L) 100 µM ATP-evoked currents in microglia at 0 mV with K^+^ and Cs^+^ in pipette. (M) Basal current in ARH microglia response to repeatedly puffed ATP (100 µM) during perfusion of TPA (50 µM). (N) Representative resting membrane potential of ARH microglia response to TPA puff (50 µM). (O) Representative basal THIK-1 current of ARH microglia recorded at 0 mV after chow or HFD exposure for 1 week in male Tmem119-CreER/Rosa26-LSL-tdTomato mice (left). Statistics for THIK-1 current of microglia following chow or HFD exposure. N = 11 (chow) or 8 (HFD) microglia from 3 different mice in each group (right). \*\**p* < 0.01. (P) Typical current response of ARH microglia to voltage steps in 20 mV increments from -104 to +56 mV (from a holding potential of -34 mV) under 1 mM (n = 7) or 5 mM (n = 7) glucose. \**p* < 0.05. (Q) Representative THIK-1 current responses to glucose fluctuation (1 → 5 mM or 5 →1 mM) in ARH microglia. (R) Statistics for THIK-1 current of microglia in the ARH, under glucose exposure from 1 → 5 mM (n = 7) or 5 →1 mM (n = 8). Significant differences between groups are shown as \*\**p* < 0.01 and \*\*\**p* < 0.001. (S) Representative THIK-1 current (top) and resting membrane potential (bottom) responses to glucose fluctuation (1 → 5 mM) under 50 µM TPA perfusion in ARH microglia. (T) THIK-1 current (left, n = 7) and resting membrane potential (right, n = 8) of microglia in the ARH, under TPA perfusion and glucose exposure from 1 → 5 mM. No differences between groups are shown as ns. Data are mean ± SEM in (P), mean ± SEM with individual data points in (D, O), and individual data points in (F, R, T). Two-sided unpaired t-test was used in (D, O); two-sided paired t-test was used in (F, R, T); two-way ANOVA analysis was used in (P).

To identify ion channel that may mediate this glucose-sensing property, we analyzed single-nucleus RNA-Seq data from mouse ARH tissue^37^. Among the top ion channels detected in ARH microglia is *KCNK13* which encodes the two-pore domain K^+^ channel THIK-1 (Figure 1G and 1H). Notably, THIK-1 has been reported to regulate microglial ramification and surveillance by controlling the microglial resting membrane potential^32^. To further define the functional contribution of THIK-1, we examined ATP-evoked responses in microglia. Perfusion of 2 mM ATP onto ARH-containing slices elicited an outwardly rectifying membrane current in microglia (Figure 1I). In parallel, puff application of 100 µM ATP hyperpolarized ARH microglia (Figure 1J). Similarly, puffing 100 µM ATP also induced an outwardly rectifying current that was abolished when intracellular K⁺ was replaced with Cs⁺ (Figure 1K and 1L), indicating that the ATP-evoked current is carried by K⁺ ions. Given that THIK-1 channel is known to be activated by ATP, we investigated whether they contribute to K^+^ conductance of microglia. Perfusion of 50 µM TPA, a specific THIK-1 channel blocker, suppressed the ATP-induced current and eliminated the basal current attributable to the tonically active THIK-1 channel (Figure 1M and S1A). In line with this loss of tonic K⁺ conductance, TPA application depolarized ARH microglia (Figure 1N), confirming that THIK-1 contributes to the maintenance of the microglial resting membrane potential^32^. In addition, we recorded currents in response to voltage steps away from the resting potential and observed no evidence of voltage-gated channel activity in ARH microglia (Figure S1B). Therefore, these results indicated that ARH microglia express functional THIK-1 channel, and that THIK-1 is a key regulator of microglial resting membrane potential.

To determine whether THIK-1 contributes to the microglial resting membrane potential difference between chow- and HFD-fed states, we examined the basal THIK-1 current in ARH microglia under the voltage-clamp mode in Tmem119-CreER/Rosa26-LSL-tdTomato mice fed with chow or HFD for 1 week. We found that the one-week HFD feeding significantly increased the basal THIK-1 current in ARH microglia (Figure 1O). Given that THIK-1 carries an outward K⁺ current, this elevation provides a mechanistic explanation for the microglial hyperpolarization observed after HFD exposure. To further test whether extracellular glucose modulates THIK-1 activity, we plotted the current–voltage relationship of THIK-1 in ARH microglia under 1 mM and 5 mM glucose. Increasing extracellular glucose from 1 mM to 5 mM significantly enhanced THIK-1 current (Figure 1P). Consistently, the basal THIK-1 current was also significantly elevated when glucose was raised from 1 mM to 5 mM and was markedly reduced when glucose was lowered from 5 mM to 1 mM (Figure 1Q and 1R). Furthermore, to establish the functional role of THIK-1 in this glucose-evoked response of ARH microglia, perfusion of TPA during the 1 → 5 mM glucose fluctuation protocol abolished both the increase in basal THIK-1 current and the associated hyperpolarization of microglia (Figure 1S and 1T). Together, these results demonstrated that THIK-1 channel mediate the glucose-sensing properties of ARH microglia and contribute to microglial hyperpolarization in the context of overnutrition.

### TPA suppresses food intake and body weight through microglia

The ARH is a critical regulator of energy homeostasis^38,39^. To determine whether THIK-1 in ARH microglia contributes to this regulation, we assessed the metabolic effects of pharmacologically inhibiting THIK-1 channel. A single intraperitoneal (i.p.) injection of the THIK-1 channel blocker TPA (2 mg/kg) in chow-fed wild-type (WT) lean male mice significantly suppressed food intake (Figure 2A and 2B). Importantly, TPA administration did not cause kaolin intake, indicating that nausea does not contribute to the hypophagic effect (Figure 2B). In addition, we used TSE metabolic cages to measure energy expenditure and locomotor activity. Acute TPA treatment did not alter energy expenditure, respiratory exchange ratio, or physical activity (Figure S2A). Furthermore, TPA does not induce anxiety-like behaviors in the open field test (Figure S2B and S2C). We next investigated whether sustained THIK-1 inhibition produces longer-term metabolic effects. Chronic daily TPA administration (2 mg/kg/day, i.p. for 7 days) in chow-fed WT lean male mice significantly reduced cumulative food intake and decreased body weight gain relative to saline controls (Figure 2C and 2D). Thus, these results demonstrated that both acute and chronic TPA treatments robustly suppress food intake without causing sickness or altering energy expenditure.

**Figure 2.**
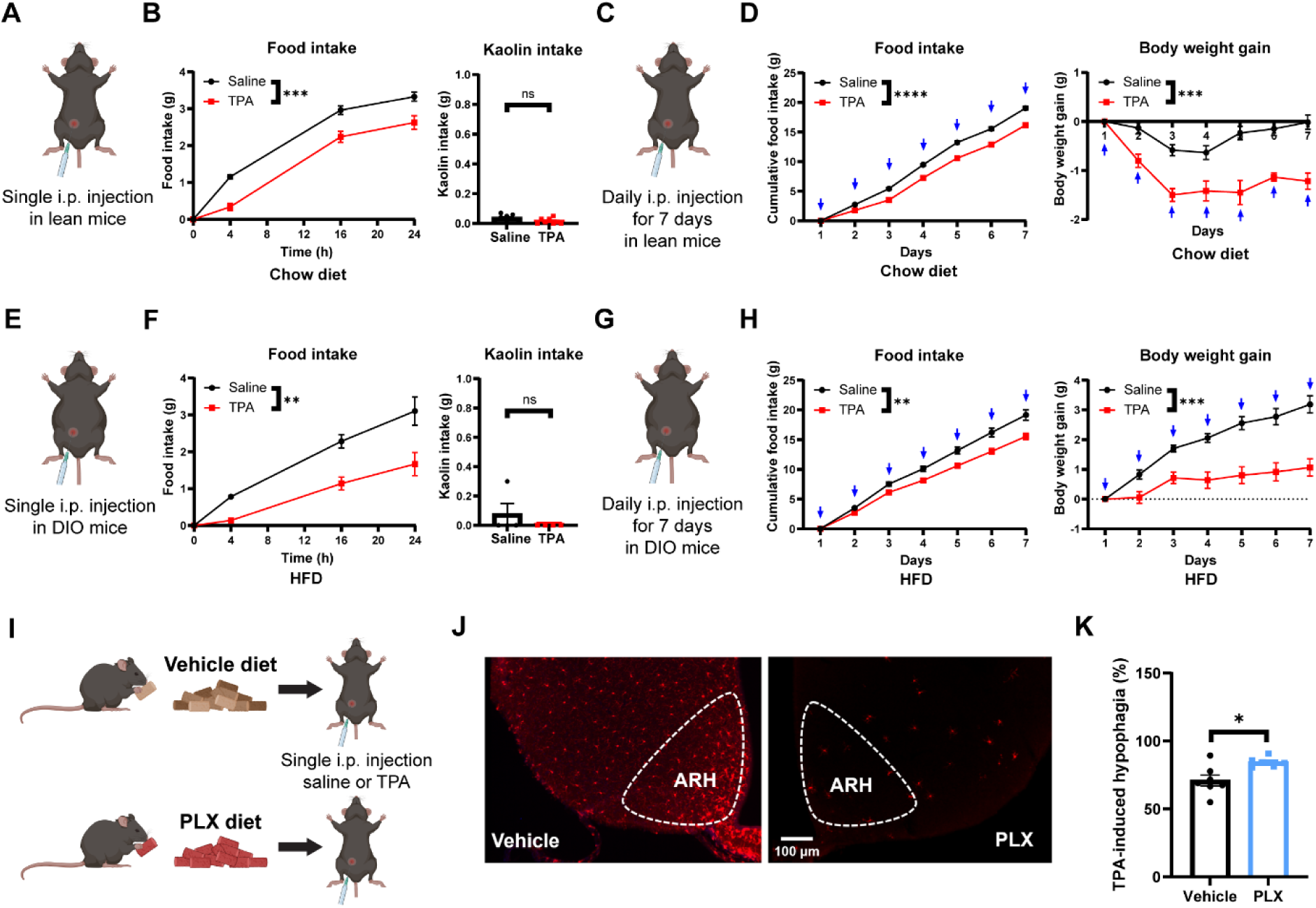
TPA suppresses food intake and body weight through microglia. (A) Schematic of the saline or TPA (2 mg/kg, i.p.) treatment in male WT lean mice. (B) Cumulative chow diet food intake and kaolin intake (4 h) of male WT lean mice following injection of either saline (n = 8) or TPA (2 mg/kg, i.p., n = 8). Significant differences between groups are shown as \*\*\**p* < 0.001. No differences between groups are shown as ns. (C) Schematic of the daily saline or TPA (2 mg/kg, i.p.) treatment in male WT lean mice. (D) Cumulative chow diet food intake and body weight gain of male WT lean mice following daily injection of either saline (n = 6) or TPA (2 mg/kg, i.p., n = 6). Significant differences between groups are shown as \*\*\**p* < 0.001 and \*\*\*\**p* < 0.0001. (E) Schematic of the saline or TPA (2 mg/kg, i.p.) treatment in male WT DIO mice. (F) Cumulative HFD food intake and kaolin intake (4 h) of male WT DIO mice following injection of either saline (n = 4) or TPA (2 mg/kg, i.p., n = 4). Significant differences between groups are shown as \*\**p* < 0.01. No differences between groups are shown as ns. (G) Schematic of the daily saline or TPA (2 mg/kg, i.p.) treatment in male WT DIO mice. (H) Cumulative HFD food intake and body weight gain of male WT DIO mice following daily injection of either saline (n = 7) or TPA (2 mg/kg, i.p., n = 8). Significant differences between groups are shown as \*\*\**p* < 0.001 and \*\**p* < 0.01. (I) Schematic of the saline or TPA (2 mg/kg, i.p.) treatment in male WT mice under vehicle diet or PLX5622 diet exposure for 1 week. (J) Representative microscopic images showing microglial Iba1 immunostaining (red) in the ARH of male WT mice under vehicle diet or PLX5622 diet exposure for 1 week. (K) Results of TPA-induced hypophagia percentage (TPA treated 24 h food intake/saline treated 24 h food intake) in male WT vehicle diet (n = 7) or PLX diet (n = 6) treated mice. Significant differences between groups are shown as \**p* < 0.05. Data are mean ± SEM in (B, D, F, H), mean ± SEM with individual data points in (K). Two-sided unpaired t-test was used in (B, F, K); two-way ANOVA analysis was used in (B, D, F, H).

To extend these findings to an obesity context, we tested TPA in a separate cohort of male WT diet-induced obese (DIO) male mice. Consistently, a single i.p. injection of TPA (2 mg/kg) produced a significant reduction in food intake without affecting kaolin consumption (Figure 2E and 2F). To determine whether this effect persists with repeated treatment, we administered TPA daily (2 mg/kg/day, i.p. for 7 days) into DIO mice. Chronic TPA treatment resulted in a sustained and significant suppression of food intake and reduced body-weight gain (Figure 2G and 2H). Together, these results demonstrated that prolonged THIK-1 inhibition attenuates hyperphagia and body-weight gain in obese mice.

Given the enriched expression of THIK-1 in microglia, we next investigated whether microglia are required for TPA-induced feeding suppression. To this end, we depleted microglia using the colony-stimulating factor 1 receptor inhibitor PLX5622. WT mice were fed chow diet containing vehicle or PLX5622 (1.2 g/kg) for 7 days, followed by a single i.p. injection of saline or TPA (Figure 2I). As expected, oral PLX5622 treatment produced near-complete microglial elimination in the ARH within 7 days, as indicated by the loss of Iba1⁺ microglia (Figure 2J). Consistent with our results, TPA induced robust hypophagia in vehicle-treated mice, whereas this effect was markedly blunted in PLX-treated mice (Figure 2K). These results indicated that microglial depletion abolishes TPA-induced anorexia, supporting a microglia-dependent mechanism through which THIK-1 inhibition regulates energy balance.

### TPA inhibits AgRP neurons through microglia-mediated mechanism

Neurons in the ARH that co-express neuropeptide Y (NPY) and AgRP define the AgRP neuron population, and activation of these AgRP neurons promotes feeding^40,41^. In parallel, recent studies have shown that microglia–neuron interactions critically regulate fundamental neural processes, including synaptic pruning, axonal remodeling, and neurogenesis^42^. To explore how inhibition of the microglial THIK-1 channel leads to feeding suppression, we performed brain-slice electrophysiological recordings to monitor microglia–AgRP neuron interaction in the ARH. In AgRP-IRES-Cre/Rosa26-LSL-tdTomato male mice (AgRP neurons are labeled with tdTomato in this mouse line), TPA perfusion (50 µM) significantly hyperpolarized AgRP neurons and reduced their firing frequency, indicating that TPA suppresses AgRP neuronal activity (Figure 3A-3C). However, direct puff application of TPA (5 s, 50 µM) onto AgRP neurons failed to alter either firing frequency or resting membrane potential (Figure 3D-3F), suggesting that TPA does not act directly on AgRP neurons. Supporting this interpretation, our single-nucleus RNA-seq data revealed weak THIK-1 expression in AgRP neurons (Figure S3A), and TPA perfusion did not affect basal currents in these neurons (Figure S3B), consistent with a lack of functional THIK-1 channel in AgRP neurons. Together, these findings indicated that TPA inhibits AgRP neuron activity through an indirect mechanism. Importantly, the TPA-induced hyperpolarization persisted even in the presence of a cocktail of synaptic blockers, 1 μM TTX (a sodium channel inhibitor), 20 μM DNQX (an AMPA glutamatergic receptor inhibitor), 50 μM D-AP5 (an NMDA glutamatergic receptor inhibitor) and 50 μM bicuculline (a GABAA receptor inhibitor; Figures 3G-3I). This persistence demonstrated that the inhibitory effects of TPA on AgRP neurons are independent of network circuitry or presynaptic input.

**Figure 3.**
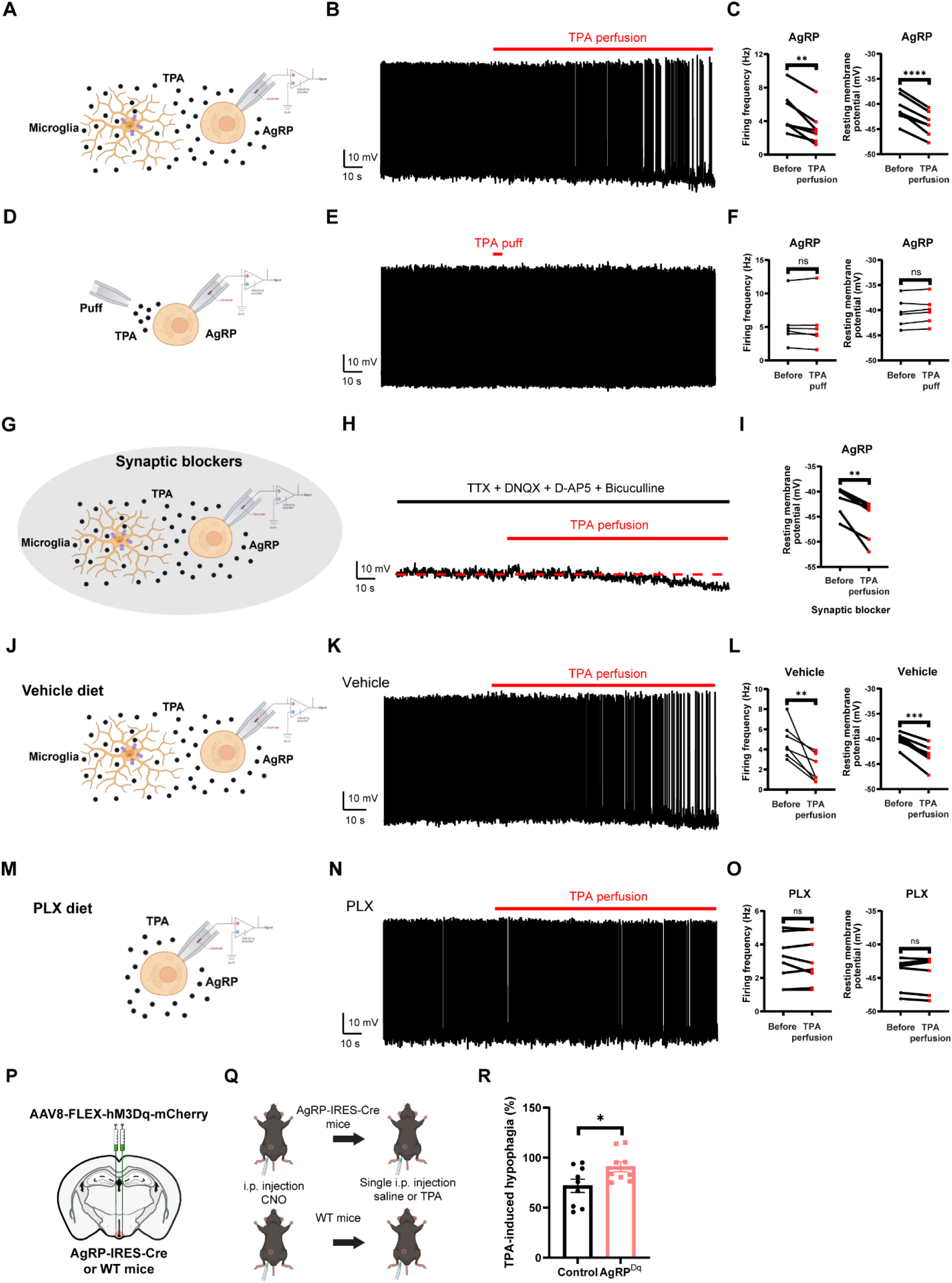
Microglia-mediated inhibition of AgRP neurons by TPA leads to feeding suppression. (A) Schematic of the AgRP neuron recording in the ARH after TPA perfusion to the brain slice. (B) A representative current clamp trace showing effects of TPA (50 µM, perfusion) on AgRP neurons in AgRP-IRES-Cre/Rosa26-LSL-tdTomato male mice. (C) Firing frequency and resting membrane potential changes of AgRP neurons after TPA perfusion; n = 7 neurons from three different mice. Significant differences between groups are shown as \*\**p* < 0.01 and \*\*\*\**p* < 0.0001. (D) Schematic of the AgRP neuron recording in the ARH after TPA puff to the AgRP neuron. (E) A representative current clamp trace showing effects of TPA (50 µM, puff for 5 s) on AgRP neurons in AgRP-IRES-Cre/Rosa26-LSL-tdTomato male mice. (F) Firing frequency and resting membrane potential changes of AgRP neurons after TPA puff; n = 6 neurons from three different mice. No differences between groups are shown as ns. (G) Schematic of the AgRP neuron recording plus the synaptic blockers TTX, DNQX, D-AP5 and bicuculline after TPA perfusion to the brain slice. (H) A representative current clamp trace showing effects of TPA (50 µM, perfusion) plus the synaptic blockers on AgRP neurons in AgRP-IRES-Cre/Rosa26-LSL-tdTomato male mice. (I) Resting membrane potential changes of AgRP neurons after TPA perfusion; n = 6 neurons from three different mice. Significant differences between groups are shown as \*\**p* < 0.01. (J) Schematic of the AgRP neuron recording in the ARH after TPA perfusion to the brain slice under vehicle diet treatment. (K) A representative current clamp trace showing effects of TPA (50 µM, perfusion) on AgRP neurons in AgRP-IRES-Cre/Rosa26-LSL-tdTomato male mice treated with vehicle diet. (L) Firing frequency and resting membrane potential changes of AgRP neurons after TPA perfusion; n = 7 neurons from three different mice. Significant differences between groups are shown as \*\**p* < 0.01 and \*\*\**p* < 0.001. (M) Schematic of the AgRP neuron recording in the ARH after TPA perfusion to the brain slice under PLX diet treatment. (N) A representative current clamp trace showing effects of TPA (50 µM, perfusion) on AgRP neurons in AgRP-IRES-Cre/Rosa26-LSL-tdTomato male mice treated with PLX diet. (O) Firing frequency and resting membrane potential changes of AgRP neurons after TPA perfusion; n = 8 neurons from three different mice. No differences between groups are shown as ns. (P) Schematic illustration for stereotaxic injection of AAV8-FLEX-hM3Dq-mCherry into the ARH of AgRP-IRES-Cre mice (AgRP^Dq^) or WT mice (control). (Q) Schematic of the saline or TPA (2 mg/kg, i.p.) treatment in AgRP-IRES-Cre mice or WT mice after CNO (2 mg/kg, i.p.) injection. (R) Results of TPA-induced hypophagia percentage (TPA treated 24 h food intake/saline treated 24 h food intake) in control mice (n = 9) or AgRP^Dq^ mice (n = 9) after CNO (2 mg/kg, i.p.) treatment. Significant differences between groups are shown as \**p* < 0.05. Data are mean ± SEM with individual data points in (R), and individual data points in (C, F, I, L, O). Two-sided unpaired t-test was used in (R); two-sided paired t-test was used in (C, F, I, L, O).

To further examine the mechanism underlying this microglia-dependent hypophagia, we recorded AgRP neuronal activity in AgRP-IRES-Cre/Rosa26-LSL-tdTomato male mice fed with either vehicle or PLX diet for 7 days. In vehicle-fed mice, where microglia remained intact, TPA perfusion (50 µM) consistently hyperpolarized AgRP neurons and reduced their firing frequency (Figure 3J–3L). Notably, in PLX-fed mice in which microglia were absent, TPA perfusion failed to alter AgRP neuron activity (Figure 3M–3O). Taken together, these findings demonstrated that microglia are required for TPA to inhibit AgRP neurons.

To further test whether inhibition of AgRP neurons is necessary for TPA-induced hypophagia, we chemogenetically activated AgRP neurons during TPA treatment. To do this, we stereotaxically injected AAV-FLEX-hM3Dq-mCherry into the ARH of AgRP-IRES-Cre mice, enabling selective expression of hM3Dq in AgRP neurons (AgRP^Dq^ mice; Figure 3P). Activation of these neurons was achieved using clozapine N-oxide (CNO, 2 mg/kg; Figure 3Q and S4). Notably, we recapitulated the hypophagic response of TPA in WT mice pre-treatment with CNO (Control, Figure 3R). In contrast, in CNO-pre-treatment AgRP^Dq^ mice in which AgRP neurons were chemogenetically activated, TPA-induced hypophagia was blunted (Figure 3R). These results demonstrated that inhibition of AgRP neurons is required for the hypophagic effect of TPA and identify AgRP neurons as a functionally relevant downstream target mediating TPA-induced hypophagia.

### TPA-induced microglial phagocytosis of PNN leads to feeding suppression

Recent work has shown that PNNs, a specialized extracellular matrix structure, progressively assemble around AgRP neurons in the ARH during postnatal maturation^43,44^. Emerging evidence indicates that PNNs regulate feeding and obesity via AgRP neurons, and that microglia contribute to AgRP neuron maturation by modulating PNN plasticity^26,45^. To examine whether microglia–AgRP neuron interaction through PNN remodeling contribute to TPA-induced hypophagia, we used Wisteria floribunda agglutinin (WFA) staining to visualize PNNs in the ARH^46^. Chronic TPA administration in chow-fed WT lean male mice (2 mg/kg/day, i.p. for 7 days) resulted in a robust decrease in both the intensity and area of PNN staining in the ARH compared with saline-treated controls (Figure 4A–4C). To determine whether this reduction reflects enhanced microglial phagocytosis of PNN components, we quantified microglial phagocytic activity using the phagolysosomal marker CD68. Co-staining for Iba1 and CD68 revealed a marked increase in CD68-positive area per microglial cell in the ARH of TPA-treated mice relative to saline controls (Figure 4D–4F). Thus, these results showed that TPA increases microglial phagocytic activity in the ARH and supports the notion that microglia-mediated PNN remodeling may contribute to the observed reduction in PNNs.

**Figure 4.**
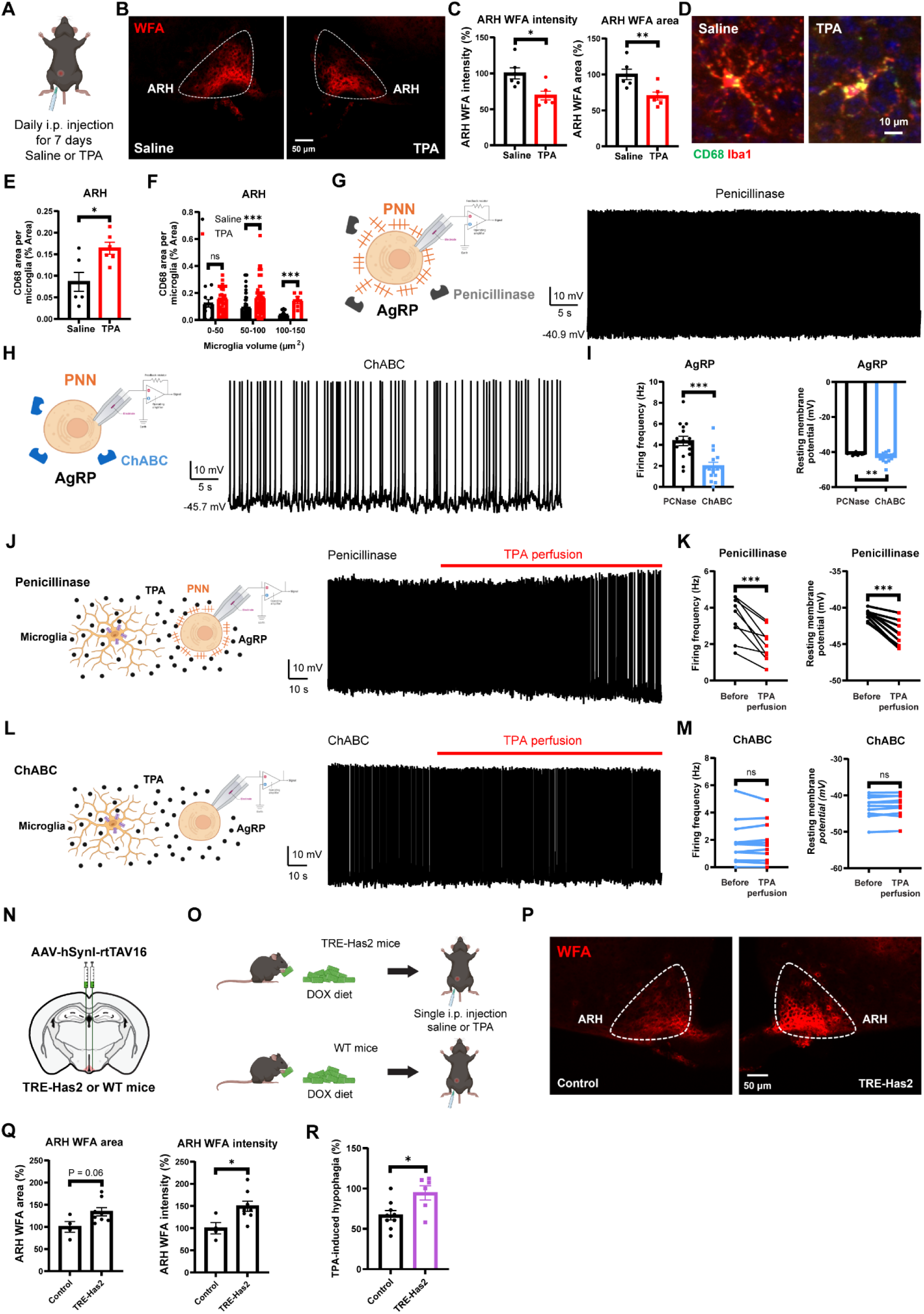
TPA-induced microglial phagocytosis of PNN leads to feeding suppression. (A) Schematic of the daily saline or TPA (2 mg/kg, i.p.) treatment in male WT lean mice. (B) Staining of PNNs using WFA in the ARH after daily saline or TPA (2 mg/kg, i.p.) treatment. (C) Statistics of the intensity and area of ARH WFA-labeled PNNs signals in saline (n = 6 mice) and TPA-treated (n = 6 mice) groups. Significant differences between groups are shown as \*\**p* < 0.01 and \**p* < 0.05. (D) Co-staining of phagolysosomal marker CD68 and microglial marker Iba1 in the ARH after daily saline or TPA (2 mg/kg, i.p.) treatment. (E) Statistics of CD68 area within the microglia (CD68 area/Iba1 area %) in saline (n = 6 mice) and TPA-treated (n = 6 mice) groups. Significant differences between groups are shown as \**p* < 0.05. (F) Statistics of CD68 area within different microglial volumes (CD68 area/Iba1 area %) in saline (n = 85 cells) and TPA-treated (n = 90 cells) groups. Significant differences between groups are shown as \*\*\**p* < 0.001. No differences between groups are shown as ns. (G-H) Schematic of the AgRP neuron recording in the ARH and representative current-clamp traces of AgRP neurons recorded under penicillinase or ChABC treatment. (I) Firing frequency and resting membrane potential of AgRP neurons in AgRP-IRES-Cre/Rosa26-LSL-tdTomato male mice under penicillinase or ChABC treatment. N = 15 (penicillinase) or 16 (ChABC) neurons from 3 different mice in each group. Significant differences between groups are shown as \*\**p* < 0.01 and \*\*\**p* < 0.001. (J-L) Schematic of the AgRP neuron recording in the ARH after TPA perfusion to the brain slice under penicillinase or ChABC treatment (left). A representative current clamp trace showing effects of TPA (50 µM, perfusion) on AgRP neurons in AgRP-IRES-Cre/Rosa26-LSL-tdTomato male mice treated with penicillinase or ChABC (right). (K) Firing frequency and resting membrane potential changes of AgRP neurons after TPA perfusion under penicillinase treatment; n = 9 neurons from three different mice. Significant differences between groups are shown as \*\*\**p* < 0.001. (M) Firing frequency and resting membrane potential changes of AgRP neurons after TPA perfusion under ChABC treatment; n = 11 neurons from three different mice. No differences between groups are shown as ns. (N) Schematic illustration for stereotaxic injection of AAV-hSynl-rtTAV16 into the ARH of TRE-Has2 mice (TRE-Has2) or WT mice (control). (O) Schematic of the saline or TPA (2 mg/kg, i.p.) treatment in TRE-Has2 mice or WT mice after DOX diet treatment. (P) Staining of PNNs using WFA in the ARH of TRE-Has2 mice or WT mice. (Q) Statistics of the intensity and area of ARH WFA-labeled PNNs signals in control (n = 4 mice) and TRE-Has2 (n = 8 mice) groups. Significant differences between groups are shown as \**p* < 0.05. (R) Results of TPA-induced hypophagia percentage (TPA treated 24 h food intake/saline treated 24 h food intake) in control (n = 9) or TRE-Has2 (n = 6) mice. Significant differences between groups are shown as \**p* < 0.05. Data are mean ± SEM with individual data points in (C, E, F, I, Q, R), and individual data points in (K, M). Two-sided unpaired t-test was used in (C, E, F, I, Q, R); two-sided paired t-test was used in (K, M).

Removal of PNNs within the ARH has been shown to inhibit AgRP neuron activity in high-fat, high-sugar (HFHS)-fed mice^45^. Building on this observation, we next examined how PNN integrity influences AgRP neuronal function in our model. We incubated ARH-containing slices from AgRP-IRES-Cre/Rosa26-LSL-tdTomato male mice with chondroitinase ABC (chABC), an enzyme that selectively digests PNNs, to mimic the PNN reduction induced by TPA and then recorded AgRP neuron activity. ChABC treatment significantly hyperpolarized AgRP neurons and reduced their firing frequency relative to penicillinase controls (Figure 4G–4I), indicating that PNN digestion suppresses AgRP neuronal activity and may contribute to TPA-induced feeding suppression.

To directly test whether PNNs are required for TPA-induced inhibition of AgRP neurons, we incubated ARH slices with either penicillinase or chABC and then applied TPA while recording AgRP neuron activity. As expected, TPA perfusion robustly inhibited AgRP neurons in penicillinase-treated slices (Figure 4J and 4K). However, in chABC-treated slices in which PNNs surrounding AgRP neurons were digested, TPA failed to induce AgRP neuron inhibition (Figure 4L and 4M). Therefore, these results suggested that PNN integrity is required for TPA-induced inhibition of AgRP neurons, identifying PNN remodeling as a key component of the microglia–AgRP neuron interaction underlying TPA’s hypophagic effects.

To further evaluate whether PNN remodeling is required for TPA-induced feeding suppression, we selectively increased PNN abundance in the ARH. To do this, we stereotaxically injected AAV-hSynI-rtTAV16 into the ARH of TRE-Has2 male mice and continuously fed them with doxycycline-containing chow diet to induce overexpression of hyaluronan synthase 2 (Has2) in the ARH (Figure 4N and 4O). WT littermate males received the same virus injections and diet and served as controls. PNNs comprise extended hyaluronan (HA) chains, and prior studies showed that Has2 upregulation increases HA levels^41^ and therefore promotes PNN formation. Consistent with this, post hoc WFA staining confirmed increased PNN area and intensity in the ARH of TRE-Has2 mice relative to WT controls (Figure 4P and 4Q). We next assessed whether enriched PNNs alter the feeding response to TPA. Notably, we recapitulated the TPA-induced feeding suppression in control group; however, this effect was markedly attenuated in TRE-Has2 mice (Figure 4R). This suggested that when PNNs are abundantly enriched around AgRP neurons, the degree of PNN reduction triggered by TPA becomes insufficient to substantially alter feeding behavior. Together, using both loss-of-function (chABC) and gain-of-function (TRE-Has2) models, we demonstrate that TPA-driven microglial phagocytosis of PNNs is a key mechanism underlying its hypophagic effect.

## DISCUSSION

In the current study, we identified a novel role for THIK-1 channel in mediating the glucose-sensing properties of ARH microglia and contributing to microglial hyperpolarization in the context of HFD-induced obesity. Notably, we demonstrated that pharmacological inhibition of THIK-1 channel by TPA robustly suppresses food intake and effectively attenuates hyperphagia and body-weight gain in obese mice. Furthermore, we found that THIK-1 inhibition in microglia reduces AgRP neuronal activity and establish microglia–AgRP neuron interaction as a key mediator of TPA’s hypophagic effect. Finally, our findings reveal that TPA-driven microglial phagocytosis of PNNs leads to AgRP inhibition, representing a central mechanism underlying its hypophagia. Together, these results support THIK-1 channel as a critical glucose sensor in ARH microglia and uncover a novel microglia-dependent pathway through which microglial THIK-1 inhibition modulates AgRP neuronal activity via PNN remodeling to regulate energy balance.

The central nervous system relies exclusively on glucose as its energy source; therefore, sensing circulating glucose levels and preventing hypoglycemia are essential for maintaining critical brain functions such as cognition, memory, and motor functions^47^. To achieve normoglycemia, specific populations of hypothalamic neurons and astrocytes sense changes in blood glucose concentrations and initiate appropriate physiological responses^47^. Microglia, the innate immune cells of the central nervous system, also sense extracellular cues and exhibit dynamic structural responses to changes in their environment^48^. Despite their well-recognized responsiveness to metabolic and inflammatory signals, whether microglia possess glucose-sensing properties has remained unknown. Here we show that elevating glucose markedly hyperpolarized ARH microglia, whereas reducing glucose depolarized them, demonstrating that microglia exhibit intrinsic glucose sensitivity. These findings provide a mechanistic explanation for the microglial hyperpolarization observed following HFD exposure. Furthermore, we identified the two-pore domain K⁺ channel THIK-1 as the glucose sensor in ARH microglia. Together, these results not only reveal a previously unrecognized glucose-sensing property of microglia but also establish THIK-1 as a central glucose sensor in microglia.

AgRP neurons are one of the major neuronal populations within the ARH that governs feeding behavior. Specifically, AgRP neurons are robustly activated during states of energy deficit to promote foraging and food intake^49–51^. Consistent with this framework, we provide multiple independent lines of evidence supporting a microglia-dependent mechanism that converges on AgRP neuron inhibition. First, depletion of microglia blunted TPA-induced feeding suppression, establishing an essential role for microglia in this process. Second, while perfusion of TPA robustly inhibited AgRP neuron activity, puff application of TPA directly onto AgRP neurons had no effect, indicating that TPA acts through an indirect pathway. Third, TPA perfusion failed to alter basal currents in AgRP neurons, suggesting that functional THIK-1 channel is absent in AgRP neurons and TPA does not directly act on AgRP neurons. Fourth, microglial deletion eliminated TPA-induced AgRP inhibition, demonstrating that intact microglia are required for this neuronal response and further supporting microglia–AgRP neuron signaling axis. Finally, functional inhibition of AgRP neurons was necessary for TPA-induced hypophagia, identifying AgRP neurons as a critical downstream effector of this response. Together, these convergent results delineate a model in which TPA suppresses feeding predominantly through inhibition of microglial THIK-1 channel, leading to secondary suppression of AgRP neuron activity.

THIK-1 channel have been identified as the principal potassium channel expressed in microglia, where their tonic activity maintains the microglial resting membrane potential and is essential for microglial ramification, surveillance, and interleukin-1β release^32^. Notably, THIK-1 expression is upregulated in microglia from multiple cortical regions of patients with Alzheimer’s disease (AD) compared with age-matched non-demented controls^52^. Similarly, elevated THIK-1 expression has been associated with inflammatory glial activation in Parkinson’s disease (PD)^53^. These observations have led to the proposal that selective inhibition of microglial THIK-1 channel, particularly at early disease stages, may provide therapeutic benefit in neurodegenerative disorders^53,54^. However, the role of THIK-1 channel in metabolic disease has remained unexplored. Here, we demonstrate that prolonged inhibition of THIK-1 attenuates hyperphagia and limits body-weight gain in obese mice. Thus, these findings extend the functional relevance of microglial THIK-1 beyond neurodegeneration and identify THIK-1 inhibition as a potential strategy for modulating microglial function in the context of obesity.

The condensed extracellular matrix structure known as PNN is composed of long glycosaminoglycans (GAGs) chains of hyaluronan (HA) arranged into a lattice-like structure surrounding a subset of neurons^55^. PNN is an essential component of the synapse, providing physical protection and ionic buffering for neurons while regulating synaptic plasticity and intracellular signaling^56^. Although PNN has long been implicated in neurobiological disorders such as schizophrenia, bipolar disorder, AD and addiction, recent findings indicate that PNNs also play important roles in energy-balance regulation^43,57^. Notably, enzymatic digestion of ARH PNN reduces food intake and body weight in DIO mice^45^. In contrast, PNN digestion within the MBH has been reported to increase food intake and body weight in mice^58^, highlighting regional heterogeneity in PNN function. Recently, emerging evidence has further revealed that microglia possess the capacity to remodel PNN structure. For example, microglia engulf PNN components surrounding AgRP neurons and microglial depletion leads to increased PNN formation^26^, suggesting that microglia play a critical role in maintaining PNN homeostasis. Within this framework, our findings identify THIK-1-dependent microglial activity as a critical regulator of PNN remodeling in the ARH. Specifically, inhibition of THIK-1 channel enhances microglial phagocytic activity, leading to the reduction of ARH PNN. Moreover, we demonstrate that PNN digestion attenuates AgRP neuronal activity and may contribute to the hypophagic effects of TPA. Thus, these results establish TPA-driven microglial phagocytosis of PNN as a key mechanism underlying its hypophagia. Therefore, our findings provide new insight into how microglia regulate PNN dynamics in the context of obesity and support a functional link between PNN remodeling and metabolic homeostasis.

It is worth noting that systemic (i.p.) administration of TPA may also influence microglial populations outside the ARH, and future studies will be required to selectively manipulate THIK-1 channel specifically in ARH microglia. Although recent advances in engineered AAV capsids have improved microglial gene delivery^59^, viral transduction of microglia remains technically challenging due to low transduction efficiency, insufficient transgene expression, and the risk of inducing immune activation. In our attempts to address this, we injected AAV11-sgKCNK13-Flex-GFP into the ARH of Tmem119-CreER/Rosa26-LSL-tdTomato mice to selectively delete THIK-1 in microglia. Despite repeated efforts, AAV expression in microglia remained undetectable, underscoring the current barriers to region-specific microglial manipulation. In parallel, it is important to consider the potential contribution of other neuronal populations within the ARH. POMC neurons represent a major neuronal subtype that exerts metabolic effects opposite to AgRP neurons. Specifically, POMC neurons are preferentially engaged during positive energy balance to suppress food intake and increase energy expenditure^60,61^. Notably, our single-nucleus RNA-seq analysis of mouse ARH tissue revealed substantial THIK-1 expression in POMC neurons, raising the possibility that TPA might also act directly on neuronal THIK-1 channel to regulate feeding behavior. Therefore, determining whether THIK-1 signaling in POMC neurons contributes to TPA-induced hypophagia needs further investigation.

In summary, we identified ARH microglia that exhibits glucose-sensing properties and establish the THIK-1 channel as a critical glucose sensor in microglia. Notably, THIK-1 inhibition by TPA elicits hypophagia through the microglia–AgRP neuron interaction, revealing the mechanism by which microglial activity shapes hypothalamic feeding circuits. Importantly, TPA-driven microglial phagocytosis of PNNs emerges as a central component of this interaction, providing mechanistic insight into how microglia remodel the synaptic microenvironment to regulate AgRP neuronal activity. Together, these findings elucidate a molecular and neurobiological framework through which microglia contribute to metabolic homeostasis and highlight microglial THIK-1 as a potential therapeutic target for treatment of diet-induced obesity.

## EXPERIMENTAL MODEL AND STUDY PARTICIPANT DETAILS

### Animals

Several mouse strains were used in the current study. AgRP-IRES-Cre mice^62^ (Jackson Lab, 012899) were crossed with WT mice to generate AgRP-IRES-Cre and WT littermates. Some AgRP-IRES-Cre mice were crossed with Rosa26-LSL-tdTomato mice to generate AgRP-IRES-Cre/Rosa26-LSL-tdTomato mice. In addition, we also crossed inducible Tmem119-CreER mice with Rosa26-LSL-tdTomato mice to generate Tmem119-CreER/Rosa26-LSL-tdTomato mice, and these mice received tamoxifen injections (0.2 mg/g, i.p., at 8-12 weeks of age) to induce tdTomato expression in microglia. For TRE-Has2 mice^63^, regular chow was provided before surgery and then a doxycycline (DOX, 600 mg/kg) chow diet (Bio-Serv, S4107) as indicated after surgery. These mice were housed in a temperature-controlled environment using a 12-hour light and 12-hour dark cycle (6 pm-6 am). Mice were individually housed at least 1 week before the study, when animals were handled daily to acclimatize the mice to human touching. Mice were weaned on a standard chow diet (Pico Lab, LabDiet, 5V5R). The mice were fed on a standard chow diet (6.5% fat, Harlan-Teklad, Madison, Wisconsin, #2920) or high fat diet (60% kcal from fat; Research Diets, D12492), unless mentioned otherwise. Food and water were provided ad libitum, unless mentioned otherwise. Care of all animals and procedures was approved by the Baylor College of Medicine Institutional Animal Care and Use Committees.

## METHOD DETAILS

### Slice electrophysiology

Tmem119-CreER/Rosa26-LSL-tdTomato mice (with 0.2 mg/g tamoxifen induction at 8-12 weeks) were used for recordings from microglia in the ARH. AgRP-IRES-Cre/Rosa26-LSL-tdTomato mice were used for recordings from AgRP neurons in the ARH. AgRP-IRES-Cre mice injected with AAV-FLEX-hM3Dq-mCherry were used for recordings from AgRP neurons in the ARH. Mice were deeply anesthetized with isoflurane and transcardially perfused with a modified ice-cold sucrose-based cutting solution (pH 7.3) containing 10 mM NaCl, 25 mM NaHCO3, 195 mM Sucrose, 5 mM Glucose, 2.5 mM KCl, 1.25 mM NaH2PO4, 2 mM Na-Pyruvate, 0.5 mM CaCl2, and 7 mM MgCl2, bubbled continuously with 95% O2 and 5% CO2. The mice were then decapitated, and the entire brain was removed and immediately submerged in the cutting solution. Slices (220 µm) were cut with a Microm HM 650V vibratome (Thermo Scientific). Four brain slices containing the ARH were obtained for each animal (bregma -1.22 mm to −1.94 mm). The slices were recovered for 30 min at 34°C and then maintained at room temperature in artificial cerebrospinal fluid (aCSF, pH 7.3) containing 126 mM NaCl, 2.5 mM KCl, 2.4 mM CaCl2, 1.2 mM NaH2PO4, 1.2 mM MgCl2, 5 mM glucose, and 21.4 mM NaHCO3 saturated with 95% O2 and 5% CO2 before recording. Slices were transferred to a recording chamber and allowed to equilibrate for at least 10 min before recording. The slices were superfused at 34°C in oxygenated aCSF at a flow rate of 1.8-2 ml/min. TdTomato-labeled microglia and tdTomato-labeled AgRP neurons in the ARH were visualized using epifluorescence and infrared–differential interference contrast (IR-DIC) imaging on an upright microscope (Eclipse FN-1, Nikon) equipped with a moveable stage (MP-285, Sutter Instrument). Patch pipettes with resistances of 3–5 MΩ were filled with intracellular solution (pH 7.3) containing 128 mM potassium gluconate, 10 mM KCl, 10 mM HEPES, 0.1 mM EGTA, 2 mM MgCl2, 0.05 mM (Na)2GTP, and 0.05 mM (Mg)ATP. Recordings were made using a MultiClamp 700B amplifier (Axon Instruments), sampled using Digidata 1440A, and analyzed offline with pClamp 10.3 software (Axon Instruments). Series resistance was monitored during the recording, and the values were generally <10 MΩ and were not compensated. The liquid junction potential was +12.5 mV and was corrected after the experiment. Data were excluded if the series resistance exceeding 20% changed during the experiment or without overshoot for action potential. Currents were amplified, filtered at 1 kHz, and digitized at 10 kHz. The current-clamp mode was engaged to test microglial resting membrane potential at the baseline of 5 mM glucose aCSF or 1 mM glucose aCSF. The values for resting membrane potential were averaged within a 1-minute bin at the 5 mM glucose or 1 mM glucose aCSF condition. We defined glucose-sensing properties by quantifying changes at least 2 mV in resting membrane potential following switches between different glucose concentrations^34^. The voltage-clamp mode was engaged to test ATP-induced THIK-1 current and basal THIK-1 current. The current-clamp mode was engaged to test AgRP neural firing frequency and resting membrane potential at the baseline and after perfusion (50 µM) or puff (5 s, 50 µM) delivery of TPA. To ensure each recorded neuron received the same amount of puff treatment, the neurons located on the surface of the slice were selected to record and the puff pipette was always put at a 100 μm horizontal and 100 μm vertical distance from the recorded neurons. The puff strength was maintained at the same level using a repeatable pressure pulse system (Picospritzer III, Parker). In some experiments, the aCSF solution containing 1 μM tetrodotoxin (TTX, Tocris), 50 μM bicuculline (Tocris), 20 μM DNQX (Tocris) and 50 μM D-AP5 (Tocris) was used to block the majority of presynaptic inputs. Each neuron was recorded for at least 1 minute at baseline and only the neurons with stable baseline were used to test the TPA treatment. The values of resting membrane potential and firing frequency were averaged in 40 seconds at each time point indicated in the figures. For the validation of CNO effect on hM3Dq-expressing AgRP neurons, neural firing frequency and resting membrane potential at the baseline and after puff delivery of CNO (5 s at 10 μM) were analyzed. Degradation of PNNs in brain slices prior to recordings was performed as previously described^64^. Briefly, brain slices were transferred and incubated in chondroitinase (ChABC)-containing ASCF (the final concentration of ChABC is 0.5 U/ml) for 45 min at 34 °C. After the incubation, slices were transferred to the recording chamber at 34°C and perfused continuously with oxygenated ACSF at a flow rate of 1.8–2.0 ml per min.

### Immunostaining

Mice were anaesthetized with volatilized isoflurane and perfused with saline followed by 10% formalin. Brain sections were cut at 30 μm and collected into five consecutive series. To assess PNN expression, one series of brain sections were incubated in biotin-labelled Wisteria floribunda agglutinin (WFA, 1:1000, L1516, Sigma-Aldrich) in PBS containing 0.25% Triton X-100 on shaker at 4°C for overnight. The sections were then washed and incubated for 2 hours at room temperature in streptavidin-Alexa Fluor 594 (1:500, S11227, Invitrogen). Signals were quantified in the neuron bodies and fibers using image J software. PNNs’ fluorescence intensity for each image was quantified using a method previously described in ImageJ by the macro plugin “Perineuronal net Intensity Program for the Standardization and Quantification of ECM Analysis” (PIPSQUEAK AI v5.3.9, Rewire Neuro, Inc.) following the method of region of interest (“ROI”)^65^. Briefly, the “Rolling Ball Radius” function of ImageJ, which removes smooth continuous background, was first used to decrease variability in background staining across the image. A ROI lacking PNNs was then created manually within four regions of the image, and the average intensity and standard deviation (SD) of each of these ROIs were determined using ImageJ. Higher value ROI was selected to represent the background. To distinguish the PNNs structure from general “loose” extracellular matrix (ECM) staining, a threshold was set at two SD above the background, and only signals above this threshold were considered PNNs and taken into quantification. All pixels below this threshold were set to “not a number.” Then, the ROI was created if PNNs enmeshed at least 2/3 of the soma. The average intensity of PNNs were recorded and calculated by PIPSQUEAK AI.

To validate microglial ablation by PLX5622, brain sections from control and PLX-treated mice were blocked (3% Normal donkey serum) for 1 hour, incubated with Rabbit anti-Iba1 antibody (1:2,000, AB178846, Abcam) on shaker at 4°C for overnight. Sections were subsequently incubated for 2 h at room temperature with donkey anti-rabbit Alexa Fluor 594 secondary antibody (1:400; A21207, Invitrogen). To evaluate whether inhibition of THIK-1 channel affects microglial phagocytosis, brain sections from control and TPA-treated mice were incubated with Rabbit anti-Iba1 and Rat anti-CD68 (1:1,000, ab53444, Abcam) on shaker at 4°C for overnight, followed by incubation for 2 h at room temperature with donkey anti-rabbit Alexa Fluor 594 and donkey anti-rat Alexa Fluor 488 secondary antibodies (1:400, A-11006, Thermo Fisher). Slides were cover-slipped and analyzed using a fluorescence microscope.

### Stereotaxic surgery

Mice were anesthetized (with 2% isoflurane) and placed in a stereotaxic instrument. Artificial eye ointment was applied to prevent corneal drying, and a heat pad was used to hold body temperature at 37°C. To specifically activate AgRP neurons, AAV8-FLEX-hM3Dq-mCherry was delivered into the ARH of AgRP-IRES-Cre mice (Addgene, #44361, titer: 5x1012 GC/ml, 0.2 µl/injection; AP: -1.7 mm; ML: ±0.2 mm; DV: -5.9 mm). To specifically overexpress PNN in ARH, TRE-Has2 mice^63^ and WT mice (male, 8 weeks of age) received AAV-hSynl-rtTAV16 virus (Addgene, Cat #102367, 1.86X10*12 GC/mL, 150ul, packaged by the Jan and Dan Duncan Neurological Research Institute’s Neuroconnectivity Core) into the ARH (AP: -1.7 mm; ML: ±0.2 mm; DV: -5.9 mm), followed by a DOX chow diet to induce the expression of rtTA. At end of experiments, all mice were perfused with 10% formalin. Brain sections were collected and cut at 30 µm, the expression of virus was checked, only those with accurate targeting were included for data analyses.

### Chemogenetics

After allowing 4 weeks for virus expression (hM3Dq, as described above), mice were individually housed for a minimum of 1 week before they were subjected to food intake measurement studies. On the day of measurement, food was removed from the cage at 12pm. CNO (2 mg/kg, i.p.) was administrated at 5:30pm, while saline or TPA (2 mg/kg, i.p.) was injected at 6pm. Pre-weighted food was returned to the cage at 6:00pm. Food intake was monitored at the time points as indicated in figures.

### Intraperitoneal injections, food intake, body weight, and energy expenditure measurement

For the acute food intake measurement in all studies, mice were fasted from 12:00pm-5:00pm. TPA was administered at 2 mg/kg body weight at 6:00pm. Pre-weighted food was returned to the cage at 6:00pm. Food intake was measured at the time points as indicated in each figure. For the daily food intake and body weight measurement in all studies, TPA was administered at 2 mg/kg body weight at 6:00pm. Pre-weighed food was provided on the day prior to the first TPA injection, and both food intake and body weight were measured daily at 6:30 pm. Energy expenditure measurements were performed in temperature-controlled (23 °C) cabinets containing 16 TSE PhenoMaster metabolic cages. Mice were acclimatized to the metabolic cages for 3 days before data collection. Data collected from day 4-7 were used for analyses and energy expenditure was analyzed using the online CalR tool^66,67^.

### Open field test

Mice received TPA (2 mg/kg) injection 0.5 h before they were subjected to the test. Mice were placed in the center of an open field box (42 cm length X 42 cm width X 30 cm height) in a room with dim light and were allowed to explore the arena for 5 minutes. All animal activity was recorded with a camera placed above the box. Time spent in the center during this period was measured (Ethovision XT software). The box was wiped clean with a paper towel soaked in 70% ethanol and dried thoroughly after each test session.

### Kaolin test

Mice were injected with TPA (2 mg/kg) at 6:00 pm, then were provided with kaolin (#23022001, Research Diets) and either regular chow diet or HFD. The amount of kaolin was recorded and calculated over a 4h period.

### Microglia depletion

Microglia were depleted by administering PLX5622, an inhibitor of the colony-stimulating factor 1 receptor, in the chow^68^. PLX5622 was purchased from Chemgood LLC and formulated in regular chow (6.5% fat, Harlan-Teklad, Madison, Wisconsin, #2920) at dose: 1.2 g/Kg of diet. Mice were placed on the PLX5622 diet for 1 week before injection of TPA. Control animals received the regular chow diet without the inhibitor.

### Bioinformatics

snRNA-seq dataset from the mouse ARH were obtained from a previously published dataset generated in our lab^37^ (GSE282955). Data were analyzed using Seurat 5.0.3^69^. Microglia-specific enrichment gene analysis was assessed using the log₂-transformed ratio of gene detection rates between microglia and other glial cell types, with a pseudo-count added. Gene expression levels were quantified from the SCT-normalized assay within the Seurat object and feature plots were generated to visualize the expression of individual genes across different cell types.

### Statistics

The sample size and statistical analysis methods were planned before our study based on the nature of experiments and our previous experience. For histology studies, the same experiment was repeated in at least 3 different mice. For electrophysiological studies, at least 6 different neurons from 3 different mice were included. The data are presented as mean ± SEM and/or as individual data points. Statistical analyses were performed using GraphPad Prism to evaluate normal distribution and variations within and among groups. Methods of statistical analyses were chosen based on the design of each experiment and are indicated in figure legends. P<0.05 was statistically significant.

## Acknowledgements

The investigators were supported by grants from the USDA/CRIS (51000-064-01S to YX), American Heart Association (26POST1562706 to Qingzhuo Liu), NIH (F32DK134121 to KMC; R01DK136479 to YX), and the Silver Endowment (to YX).

## Author contributions

Qingzhuo Liu, Hailan Liu, and YX conceived the project, experimental design and writing the manuscript. Qingzhuo Liu and Hailan Liu performed the procedures, data acquisition and analyses. JCB and Jinjing Jian performed the bioinformatics analysis. Tong Zhou, Yuxue Yang, and Meixin Sun assisted in molecular and cellular experiments. Yongxiang Li, KMC, MW, YD, JQ, FW, XL, YL, JC, XW, LX, CC, CM contributed to the generation of study mice and data discussion. Yi Zhu provided the TRE-Has2 mice line. Yongjie Yang and LT were involved in study design and data discussion.

## References

1. Stuber, G.D., Schwitzgebel, V.M., and Lüscher, C. (2025). The neurobiology of overeating. Neuron 113, 1680–1693. 10.1016/j.neuron.2025.03.010.

2. Caballero, B. (2019). Humans against Obesity: Who Will Win? Adv Nutr 10, S4–s9. 10.1093/advances/nmy055.

3. Saltiel, A.R., and Olefsky, J.M. (2017). Inflammatory mechanisms linking obesity and metabolic disease. J Clin Invest 127, 1–4. 10.1172/jci92035.

4. Henry, S.L., Barzel, B., Wood-Bradley, R.J., Burke, S.L., Head, G.A., and Armitage, J.A. (2012). Developmental origins of obesity-related hypertension. Clin Exp Pharmacol Physiol 39, 799–806. 10.1111/j.1440-1681.2011.05579.x.

5. Goossens, G.H. (2017). The Metabolic Phenotype in Obesity: Fat Mass, Body Fat Distribution, and Adipose Tissue Function. Obes Facts 10, 207–215. 10.1159/000471488.

6. Luke, A., and Cooper, R.S. (2013). Physical activity does not influence obesity risk: time to clarify the public health message. Int J Epidemiol 42, 1831–1836. 10.1093/ije/dyt159.

7. Stanley, S., Moheet, A., and Seaquist, E.R. (2019). Central Mechanisms of Glucose Sensing and Counterregulation in Defense of Hypoglycemia. Endocr Rev 40, 768–788. 10.1210/er.2018-00226.

8. Alvarsson, A., and Stanley, S.A. (2018). Remote control of glucose-sensing neurons to analyze glucose metabolism. Am J Physiol Endocrinol Metab 315, E327–e339. 10.1152/ajpendo.00469.2017.

9. Levin, B.E., Dunn-Meynell, A.A., and Routh, V.H. (1999). Brain glucose sensing and body energy homeostasis: role in obesity and diabetes. Am J Physiol 276, R1223–1231. 10.1152/ajpregu.1999.276.5.R1223.

10. Marty, N., Dallaporta, M., and Thorens, B. (2007). Brain glucose sensing, counterregulation, and energy homeostasis. Physiology (Bethesda) 22, 241–251. 10.1152/physiol.00010.2007.

11. Zhou, L., Podolsky, N., Sang, Z., Ding, Y., Fan, X., Tong, Q., Levin, B.E., and McCrimmon, R.J. (2010). The medial amygdalar nucleus: a novel glucose-sensing region that modulates the counterregulatory response to hypoglycemia. Diabetes 59, 2646–2652. 10.2337/db09-0995.

12. Lamy, C.M., Sanno, H., Labouèbe, G., Picard, A., Magnan, C., Chatton, J.Y., and Thorens, B. (2014). Hypoglycemia-activated GLUT2 neurons of the nucleus tractus solitarius stimulate vagal activity and glucagon secretion. Cell Metab 19, 527–538. 10.1016/j.cmet.2014.02.003.

13. Baraibar, A.M., Ardanaz, C.G., Mato, S., Kofuji, P., Araque, A., and Solas, M. (2025). Astrocytic Glucose Sensing Drives Synaptic Depression under Metabolic Stress. Aging Dis. 10.14336/ad.2025.0507.

14. Geller, S., Zanou, N., Lagarrigue, S., Zehnder, T., Gouelle, C., Santoro, T., Repond, C., Bezzi, P., Amati, F., Bouzier-Sore, A.K., et al. (2025). Hypothalamic Astrocytes Exhibit Glycolytic Features Making Them Prone for Glucose Sensing. Glia 73, 2253–2272. 10.1002/glia.70066.

15. Douglass, J.D., Ness, K.M., Valdearcos, M., Wyse-Jackson, A., Dorfman, M.D., Frey, J.M., Fasnacht, R.D., Santiago, O.D., Niraula, A., Banerjee, J., et al. (2023). Obesity-associated microglial inflammatory activation paradoxically improves glucose tolerance. Cell Metab 35, 1613–1629.e1618. 10.1016/j.cmet.2023.07.008.

16. Valdearcos, M., Robblee, M.M., Benjamin, D.I., Nomura, D.K., Xu, A.W., and Koliwad, S.K. (2014). Microglia dictate the impact of saturated fat consumption on hypothalamic inflammation and neuronal function. Cell Rep 9, 2124–2138. 10.1016/j.celrep.2014.11.018.

17. Salter, M.W., and Stevens, B. (2017). Microglia emerge as central players in brain disease. Nat Med 23, 1018–1027. 10.1038/nm.4397.

18. Butovsky, O., Jedrychowski, M.P., Moore, C.S., Cialic, R., Lanser, A.J., Gabriely, G., Koeglsperger, T., Dake, B., Wu, P.M., Doykan, C.E., et al. (2014). Identification of a unique TGF-β-dependent molecular and functional signature in microglia. Nat Neurosci 17, 131–143. 10.1038/nn.3599.

19. Fleshner, M. (2013). Stress-evoked sterile inflammation, danger associated molecular patterns (DAMPs), microbial associated molecular patterns (MAMPs) and the inflammasome. Brain Behav Immun 27, 1–7. 10.1016/j.bbi.2012.08.012.

20. Winkler, Z., Kuti, D., Polyák, Á., Juhász, B., Gulyás, K., Lénárt, N., Dénes, Á., Ferenczi, S., and Kovács, K.J. (2019). Hypoglycemia-activated Hypothalamic Microglia Impairs Glucose Counterregulatory Responses. Sci Rep 9, 6224. 10.1038/s41598-019-42728-3.

21. Valdearcos, M., Douglass, J.D., Robblee, M.M., Dorfman, M.D., Stifler, D.R., Bennett, M.L., Gerritse, I., Fasnacht, R., Barres, B.A., Thaler, J.P., and Koliwad, S.K. (2017). Microglial Inflammatory Signaling Orchestrates the Hypothalamic Immune Response to Dietary Excess and Mediates Obesity Susceptibility. Cell Metab 26, 185–197.e183. 10.1016/j.cmet.2017.05.015.

22. De Luca, S.N., Miller, A.A., Sominsky, L., and Spencer, S.J. (2020). Microglial regulation of satiety and cognition. J Neuroendocrinol 32, e12838. 10.1111/jne.12838.

23. Fernández-Arjona, M.D.M., León-Rodríguez, A., Grondona, J.M., and López-Ávalos, M.D. (2022). Long-term priming of hypothalamic microglia is associated with energy balance disturbances under diet-induced obesity. Glia 70, 1734–1761. 10.1002/glia.24217.

24. Reis, W.L., Yi, C.X., Gao, Y., Tschöp, M.H., and Stern, J.E. (2015). Brain innate immunity regulates hypothalamic arcuate neuronal activity and feeding behavior. Endocrinology 156, 1303–1315. 10.1210/en.2014-1849.

25. Yi, C.X., Walter, M., Gao, Y., Pitra, S., Legutko, B., Kälin, S., Layritz, C., García-Cáceres, C., Bielohuby, M., Bidlingmaier, M., et al. (2017). TNFα drives mitochondrial stress in POMC neurons in obesity. Nat Commun 8, 15143. 10.1038/ncomms15143.

26. Sun, J., Wang, X., Sun, R., Xiao, X., Wang, Y., Peng, Y., and Gao, Y. (2024). Microglia shape AgRP neuron postnatal development via regulating perineuronal net plasticity. Mol Psychiatry 29, 306–316. 10.1038/s41380-023-02326-2.

27. Barsh, G.S., and Schwartz, M.W. (2002). Genetic approaches to studying energy balance: perception and integration. Nat Rev Genet 3, 589–600. 10.1038/nrg862.

28. Thaler, J.P., Yi, C.X., Schur, E.A., Guyenet, S.J., Hwang, B.H., Dietrich, M.O., Zhao, X., Sarruf, D.A., Izgur, V., Maravilla, K.R., et al. (2012). Obesity is associated with hypothalamic injury in rodents and humans. J Clin Invest 122, 153–162. 10.1172/jci59660.

29. Kim, J.D., Yoon, N.A., Jin, S., and Diano, S. (2019). Microglial UCP2 Mediates Inflammation and Obesity Induced by High-Fat Feeding. Cell Metab 30, 952–962.e955. 10.1016/j.cmet.2019.08.010.

30. Lv, Z., Chen, L., Chen, P., Peng, H., Rong, Y., Hong, W., Zhou, Q., Li, N., Li, B., Paolicelli, R.C., and Zhan, Y. (2024). Clearance of β-amyloid and synapses by the optogenetic depolarization of microglia is complement selective. Neuron 112, 740–754.e747. 10.1016/j.neuron.2023.12.003.

31. Laprell, L., Schulze, C., Brehme, M.L., and Oertner, T.G. (2021). The role of microglia membrane potential in chemotaxis. J Neuroinflammation 18, 21. 10.1186/s12974-020-02048-0.

32. Madry, C., Kyrargyri, V., Arancibia-Cárcamo, I.L., Jolivet, R., Kohsaka, S., Bryan, R.M., and Attwell, D. (2018). Microglial Ramification, Surveillance, and Interleukin-1β Release Are Regulated by the Two-Pore Domain K(+) Channel THIK-1. Neuron 97, 299–312.e296. 10.1016/j.neuron.2017.12.002.

33. He, Y., Xu, P., Wang, C., Xia, Y., Yu, M., Yang, Y., Yu, K., Cai, X., Qu, N., Saito, K., et al. (2020). Estrogen receptor-α expressing neurons in the ventrolateral VMH regulate glucose balance. Nat Commun 11, 2165. 10.1038/s41467-020-15982-7.

34. Tu, L., Bean, J.C., He, Y., Liu, H., Yu, M., Liu, H., Zhang, N., Yin, N., Han, J., Scarcelli, N.A., et al. (2023). Anoctamin 4 channel currents activate glucose-inhibited neurons in the mouse ventromedial hypothalamus during hypoglycemia. J Clin Invest 133. 10.1172/jci163391.

35. Gruetter, R., Novotny, E.J., Boulware, S.D., Rothman, D.L., Mason, G.F., Shulman, G.I., Shulman, R.G., and Tamborlane, W.V. (1992). Direct measurement of brain glucose concentrations in humans by 13C NMR spectroscopy. Proc Natl Acad Sci U S A 89, 1109–1112. 10.1073/pnas.89.3.1109.

36. Harris, R.B.S. (2017). Appetite and Food Intake (CRC Press/Taylor & Francis 2017 by Taylor & Francis Group, LLC.). 10.1201/9781315120171.

37. 37. Jonathan, C., Jian, J., Lu, T., Liu, H., Kristine, M., Darah, A., Sanika, V., Megan, E., Cheng, J., Deng, Y., Fang, X., Geng, X., Han, J., Li, Y., Liu, H., Liu, Q., Liu, Y., Shi, Y., Tu, L., Wang, M., Xu, X., Yang, Y., Yu, M., Liu, X., Sun, M., Wang, F., Olivia, Z., Yang, Y., He, Y., Wang, C., Qi, Y., Li, H., Xu, Y. (2026). Sex-specific differences in mediobasal hypothalamus in response to nutritional states. Nature Communications (in press).

38. Vohra, M.S., Benchoula, K., Serpell, C.J., and Hwa, W.E. (2022). AgRP/NPY and POMC neurons in the arcuate nucleus and their potential role in treatment of obesity. Eur J Pharmacol 915, 174611. 10.1016/j.ejphar.2021.174611.

39. Bombassaro, B., Araujo, E.P., and Velloso, L.A. (2024). The hypothalamus as the central regulator of energy balance and its impact on current and future obesity treatments. Arch Endocrinol Metab 68, e240082. 10.20945/2359-4292-2024-0082.

40. Deem, J.D., Faber, C.L., and Morton, G.J. (2022). AgRP neurons: Regulators of feeding, energy expenditure, and behavior. Febs j 289, 2362–2381. 10.1111/febs.16176.

41. Morton, G.J., Cummings, D.E., Baskin, D.G., Barsh, G.S., and Schwartz, M.W. (2006). Central nervous system control of food intake and body weight. Nature 443, 289–295. 10.1038/nature05026.

42. Hu, Y., and Tao, W. (2024). Current perspectives on microglia-neuron communication in the central nervous system: Direct and indirect modes of interaction. J Adv Res 66, 251–265. 10.1016/j.jare.2024.01.006.

43. Mirzadeh, Z., Alonge, K.M., Cabrales, E., Herranz-Pérez, V., Scarlett, J.M., Brown, J.M., Hassouna, R., Matsen, M.E., Nguyen, H.T., Garcia-Verdugo, J.M., et al. (2019). Perineuronal Net Formation during the Critical Period for Neuronal Maturation in the Hypothalamic Arcuate Nucleus. Nat Metab 1, 212–221. 10.1038/s42255-018-0029-0.

44. Lupori, L., Totaro, V., Cornuti, S., Ciampi, L., Carrara, F., Grilli, E., Viglione, A., Tozzi, F., Putignano, E., Mazziotti, R., et al. (2023). A comprehensive atlas of perineuronal net distribution and colocalization with parvalbumin in the adult mouse brain. Cell Rep 42, 112788. 10.1016/j.celrep.2023.112788.

45. Beddows, C.A., Shi, F., Horton, A.L., Dalal, S., Zhang, P., Ling, C.C., Yong, V.W., Loh, K., Cho, E., Karagiannis, C., et al. (2024). Pathogenic hypothalamic extracellular matrix promotes metabolic disease. Nature 633, 914–922. 10.1038/s41586-024-07922-y.

46. Woo, A.M., Fleischel, E.J., Patel, D.C., and Sontheimer, H. (2025). Contribution of perineuronal nets to hyperexcitability in pilocarpine-induced status epilepticus. Epilepsia 66, 3528–3543. 10.1111/epi.18489.

47. Choi, J.H., and Kim, M.S. (2022). Homeostatic Regulation of Glucose Metabolism by the Central Nervous System. Endocrinol Metab (Seoul) 37, 9–25. 10.3803/EnM.2021.1364.

48. Umpierre, A.D., and Wu, L.J. (2021). How microglia sense and regulate neuronal activity. Glia 69, 1637–1653. 10.1002/glia.23961.

49. Atasoy, D., Betley, J.N., Su, H.H., and Sternson, S.M. (2012). Deconstruction of a neural circuit for hunger. Nature 488, 172–177. 10.1038/nature11270.

50. Burnett, C.J., Li, C., Webber, E., Tsaousidou, E., Xue, S.Y., Brüning, J.C., and Krashes, M.J. (2016). Hunger-Driven Motivational State Competition. Neuron 92, 187–201. 10.1016/j.neuron.2016.08.032.

51. Alcantara, I.C., Tapia, A.P.M., Aponte, Y., and Krashes, M.J. (2022). Acts of appetite: neural circuits governing the appetitive, consummatory, and terminating phases of feeding. Nat Metab 4, 836–847. 10.1038/s42255-022-00611-y.

52. Ossola, B., Rifat, A., Rowland, A., Hunter, H., Drinkall, S., Bender, C., Hamlischer, M., Teall, M., Burley, R., Barker, D.F., et al. (2023). Characterisation of C101248: A novel selective THIK-1 channel inhibitor for the modulation of microglial NLRP3-inflammasome. Neuropharmacology 224, 109330. 10.1016/j.neuropharm.2022.109330.

53. Tang, H., Sun, Y., Fachim, H.A., Cheung, T.K.D., Reynolds, G.P., and Harte, M.K. (2023). Elevated Expression of Two Pore Potassium Channel THIK-1 in Alzheimer’s Disease: An Inflammatory Mechanism. J Alzheimers Dis 95, 1757–1769. 10.3233/jad-230616.

54. Bürli, R.W., Doyle, K.J., Dickson, L., Rowland, A., Matthews, K., Stott, A.J., Teall, M., Ossola, B., Russell, S.G., Harvey, J.R.M., et al. (2024). Discovery of CVN293, a Brain Permeable KCNK13 (THIK-1) Inhibitor Suitable for Clinical Assessment. ACS Med Chem Lett 15, 646–652. 10.1021/acsmedchemlett.4c00035.

55. Fawcett, J.W., Oohashi, T., and Pizzorusso, T. (2019). The roles of perineuronal nets and the perinodal extracellular matrix in neuronal function. Nat Rev Neurosci 20, 451–465. 10.1038/s41583-019-0196-3.

56. Testa, D., Prochiantz, A., and Di Nardo, A.A. (2019). Perineuronal nets in brain physiology and disease. Semin Cell Dev Biol 89, 125–135. 10.1016/j.semcdb.2018.09.011.

57. Alonge, K.M., Mirzadeh, Z., Scarlett, J.M., Logsdon, A.F., Brown, J.M., Cabrales, E., Chan, C.K., Kaiyala, K.J., Bentsen, M.A., Banks, W.A., et al. (2020). Hypothalamic perineuronal net assembly is required for sustained diabetes remission induced by fibroblast growth factor 1 in rats. Nat Metab 2, 1025–1033. 10.1038/s42255-020-00275-6.

58. Kohnke, S., Buller, S., Nuzzaci, D., Ridley, K., Lam, B., Pivonkova, H., Bentsen, M.A., Alonge, K.M., Zhao, C., Tadross, J., et al. (2021). Nutritional regulation of oligodendrocyte differentiation regulates perineuronal net remodeling in the median eminence. Cell Rep 36, 109362. 10.1016/j.celrep.2021.109362.

59. Lin, R., Zhou, Y., Yan, T., Wang, R., Li, H., Wu, Z., Zhang, X., Zhou, X., Zhao, F., Zhang, L., et al. (2022). Directed evolution of adeno-associated virus for efficient gene delivery to microglia. Nat Methods 19, 976–985. 10.1038/s41592-022-01547-7.

60. Mercer, A.J., Hentges, S.T., Meshul, C.K., and Low, M.J. (2013). Unraveling the central proopiomelanocortin neural circuits. Front Neurosci 7, 19. 10.3389/fnins.2013.00019.

61. De Solis, A.J., Del Río-Martín, A., Radermacher, J., Chen, W., Steuernagel, L., Bauder, C.A., Eggersmann, F.R., Morgan, D.A., Cremer, A.L., Sué, M., et al. (2024). Reciprocal activity of AgRP and POMC neurons governs coordinated control of feeding and metabolism. Nat Metab 6, 473–493. 10.1038/s42255-024-00987-z.

62. Tong, Q., Ye, C.P., Jones, J.E., Elmquist, J.K., and Lowell, B.B. (2008). Synaptic release of GABA by AgRP neurons is required for normal regulation of energy balance. Nat Neurosci 11, 998–1000. 10.1038/nn.2167.

63. Chen, X., Wang, Y., Li, H., Deng, Y., Giang, C., Song, A., Liu, Y., Wang, Q.A., and Zhu, Y. (2024). Hyaluronan Mediates Cold-Induced Adipose Tissue Beiging. Cells 13. 10.3390/cells13151233.

64. Yu, K., He, Y., Hyseni, I., Pei, Z., Yang, Y., Xu, P., Cai, X., Liu, H., Qu, N., Liu, H., et al. (2020). 17β-estradiol promotes acute refeeding in hungry mice via membrane-initiated ERα signaling. Mol Metab 42, 101053. 10.1016/j.molmet.2020.101053.

65. Slaker, M.L., Harkness, J.H., and Sorg, B.A. (2016). A standardized and automated method of perineuronal net analysis using Wisteria floribunda agglutinin staining intensity. IBRO Rep 1, 54–60. 10.1016/j.ibror.2016.10.001.

66. Mina, A.I., LeClair, R.A., LeClair, K.B., Cohen, D.E., Lantier, L., and Banks, A.S. (2018). CalR: A Web-Based Analysis Tool for Indirect Calorimetry Experiments. Cell Metab 28, 656–666.e651. 10.1016/j.cmet.2018.06.019.

67. Müller, T.D., Klingenspor, M., and Tschöp, M.H. (2021). Revisiting energy expenditure: how to correct mouse metabolic rate for body mass. Nat Metab 3, 1134–1136. 10.1038/s42255-021-00451-2.

68. Elmore, M.R., Najafi, A.R., Koike, M.A., Dagher, N.N., Spangenberg, E.E., Rice, R.A., Kitazawa, M., Matusow, B., Nguyen, H., West, B.L., and Green, K.N. (2014). Colony-stimulating factor 1 receptor signaling is necessary for microglia viability, unmasking a microglia progenitor cell in the adult brain. Neuron 82, 380–397. 10.1016/j.neuron.2014.02.040.

69. Hao, Y., Stuart, T., Kowalski, M.H., Choudhary, S., Hoffman, P., Hartman, A., Srivastava, A., Molla, G., Madad, S., Fernandez-Granda, C., and Satija, R. (2024). Dictionary learning for integrative, multimodal and scalable single-cell analysis. Nat Biotechnol 42, 293–304. 10.1038/s41587-023-01767-y.

