## Supplementary data for "THIK-1 channel mediates microglial glucose sensing and modulates AgRP neurons"

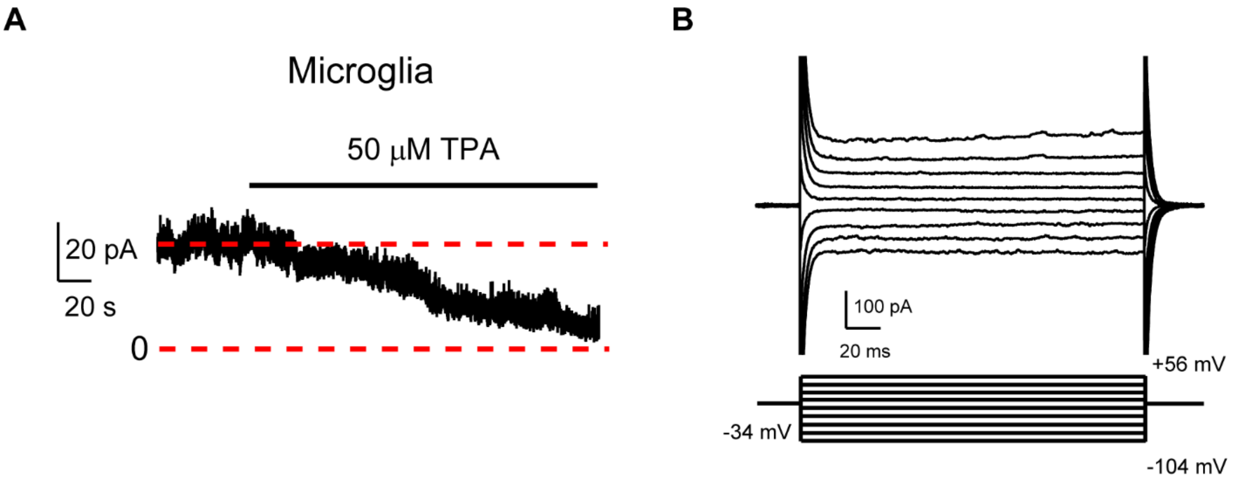


**Supplementary Figure 1. THIK-1 channel current and I-V relation in microglia.** (A) Representative THIK-1 current responses to 50 μM TPA perfusion in ARH microglia. (B) Typical current response of ARH microglia to voltage steps in 20 mV increments from -104 to +56 mV (from a holding potential of -34 mV).

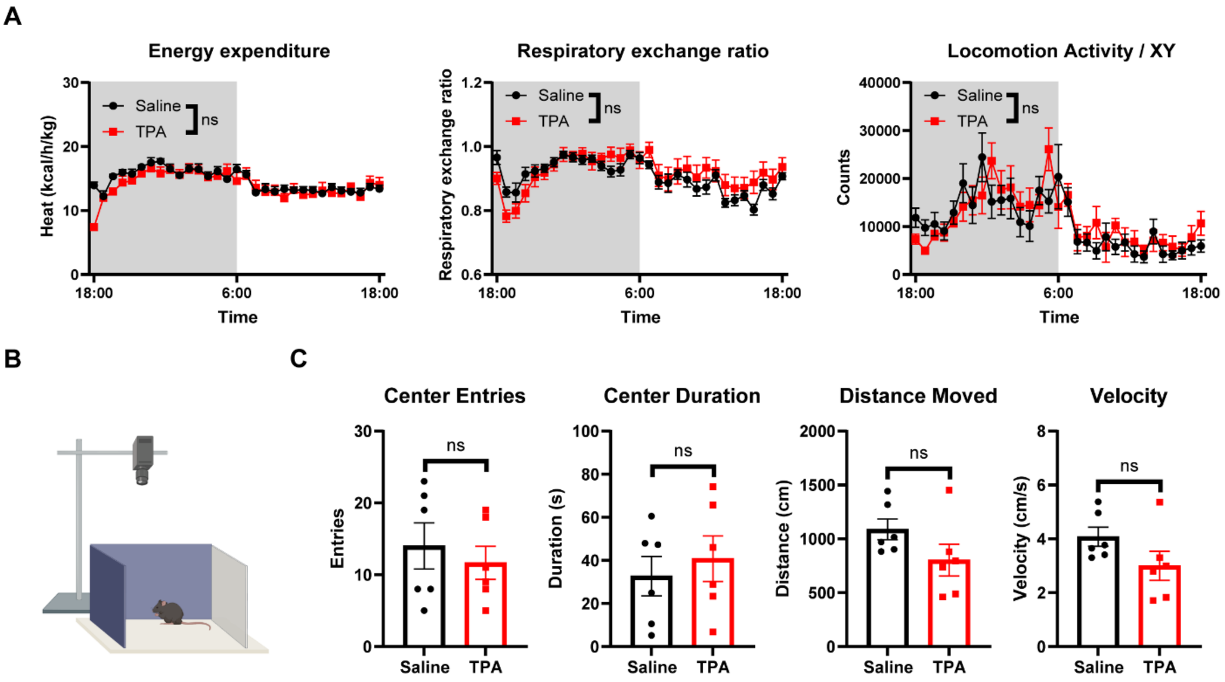


**Supplementary Figure 2. TPA has no effect on energy expenditure and does not elicit anxiety-like behavior.** (A) Energy expenditure (left), respiratory exchange ratio (middle), and locomotor activity (right) during 24 h of male WT mice following injection of either saline (n = 7) or TPA (2 mg/kg, i.p., n = 8). No differences between groups are shown as ns. (B) Schematic of open field test. (C) Number of entries into the center, time spent in the center, travel distance, and velocity during open field test in male WT mice receiving i.p. injections of saline (n = 6) or TPA (2 mg/kg, n = 6). No differences between groups are shown as ns. Data are mean ± SEM in (A). Data are mean ± SEM with individual data points in (C). Two-way ANOVA analysis was used in (A).Two-sided unpaired t-test was used in (C).

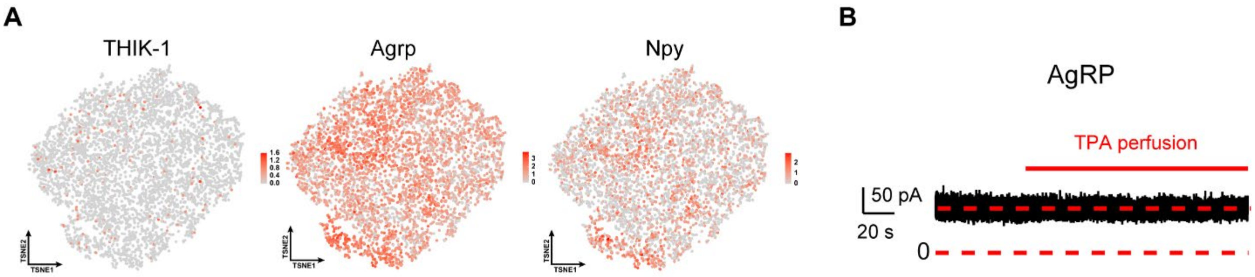


**Supplementary Figure 3. No functional THIK-1 channel in AgRP neurons.** (A) Feature plot showing THIK-1, AgRP, and NPY expression in AgRP neurons. (B) Representative basic current responses to 50 μM TPA perfusion in AgRP neurons.

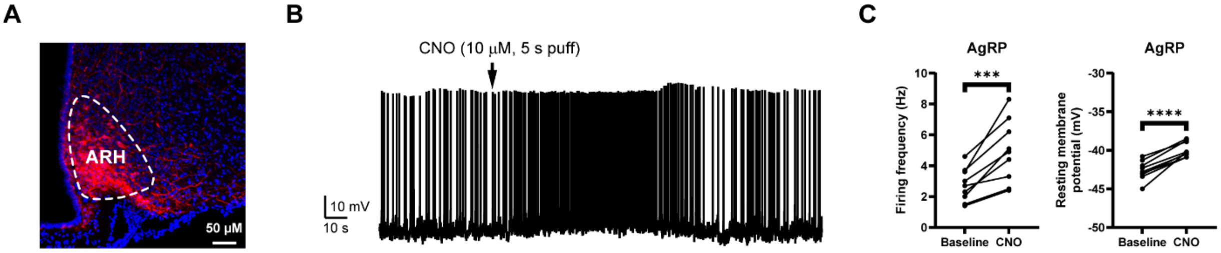


**Supplementary Figure 4. Validation of AAV8-FLEX-hM3Dq-mCherry in the ARH.** (A) Representative image illustrating the expression of mCherry in the ARH of AgRP-IRES-Cre mice. (B) A representative current-clamp traces of the recorded AgRP neuron expressing hM3Dq-mCherry in response to 10 μM CNO (puff for 5 s). (C) Firing frequency and resting membrane potential of AgRP neuron expressing hM3Dq-mCherry during the baseline and 10 μM CNO treatment. N = 9 neurons from 3 different mice. Significant differences between groups are shown as ****$p$ < 0.0001 and ***$p$ < 0.001. Data are individual data points in (C). Two-sided paired t-test was used in (C).
